# Multiplexable fluorescence lifetime imaging (FLIM) probes for Abl and Src-family kinases

**DOI:** 10.1101/655407

**Authors:** Nur P. Damayanti, Sampreeti Jena, Jackie Tan, Joseph M. K. Irudayaraj, L. L. Parker

## Abstract

Most commonly employed strategies to map kinase activities in live cells require expression of genetically-encoded proteins (e.g. FRET sensors). In this work, we describe development and preliminary application of a set of cell-penetrating, fluorophore labelled peptide substrates for fluorescence lifetime imaging (FLIM) of Abl and Src-family kinase activities. These probes do not rely on FRET pairs or genetically-encoded protein expression. We also demonstrate image-by-image and pixel-by-pixel quantification of probe phosphorylation ratio, suggesting that this strategy will be useful for detailed mapping of single cell and subcellular kinase activity in live cells.

---

Protein kinases control many fundamental aspects of cell function, including the cell cycle, DNA repair, and response to exogenous stimuli. Despite the biological importance of their dynamics and subcellular localization, few tools exist for monitoring kinase activities in cells in real time. Genetically-encoded Förster resonance energy transfer (FRET) protein sensors can accomplish single cell, dynamic imaging, however these are difficult to multiplex since two fluorophores (often with relatively broad excitation and emission spectra) are required per sensor, so spectral overlap/bleed through between sensors becomes a significant problem. We previously reported that a fluorophore labelled, cell-penetrating peptide substrate for Abl kinase exhibited longer fluorescence lifetime upon phosphorylation and apparent interaction of the phosphopeptide product with SH2 domains.^1^ This novel strategy was capable of detecting Abl kinase activity and inhibition in live cells, however it was not clear if the approach would be applicable to other kinases. We recently extended this approach to monitor apparent Akt, VEGFR2 and FAK activation in a range of cell models using 5-FAM, a fluorescein-based fluorophore label.^2, 3^ In this current report, we demonstrate that this strategy can be applied to the Src family and to additional fluorophore scaffolds (DyLight 488, a fluorescein-like scaffold, and DyLight 550, a cyanine/Cy3-like scaffold). In addition, the FLIM probes were shown to be amenable to multiplexed analysis with multi-color labelling and imaging. Time lapse imaging was employed to investigate activity of two kinases simultaneously after EGF stimulation and inactivation by an inhibitor. The proof of concept was further validated by testing of suitable negative and positive control probes. Moreover, we show for the first time that the lifetime data in each cell or even each pixel can be used to extract fractional contributions from “unphosphorylated” and “phosphorylated” probe populations, paving the way to more detailed mapping of kinase activity and phosphorylation dynamics within cell populations and throughout the interior of a live cell.

We started by exploring the generality of the fluorescence lifetime shift upon phosphorylation of the peptide probe, using DyLight 488 and DyLight 550. Two substrates were chosen: Abltide (Abl kinase substrate, as we used in previous work)^1^ and SFAStide-A (a Src-family kinase substrate previously developed by the Parker laboratory)^5^ and conjugated with TAT to create cell-penetrating probes. Peptides were synthesized with the sequences shown in Table 1, labelled on the central cysteine residue by maleimide chemistry, and purified by preparative HPLC (as previously described^1^). The corresponding phosphorylated and Y→F mutant probes were also synthesized as positive and negative controls, respectively.

**Table 1.**
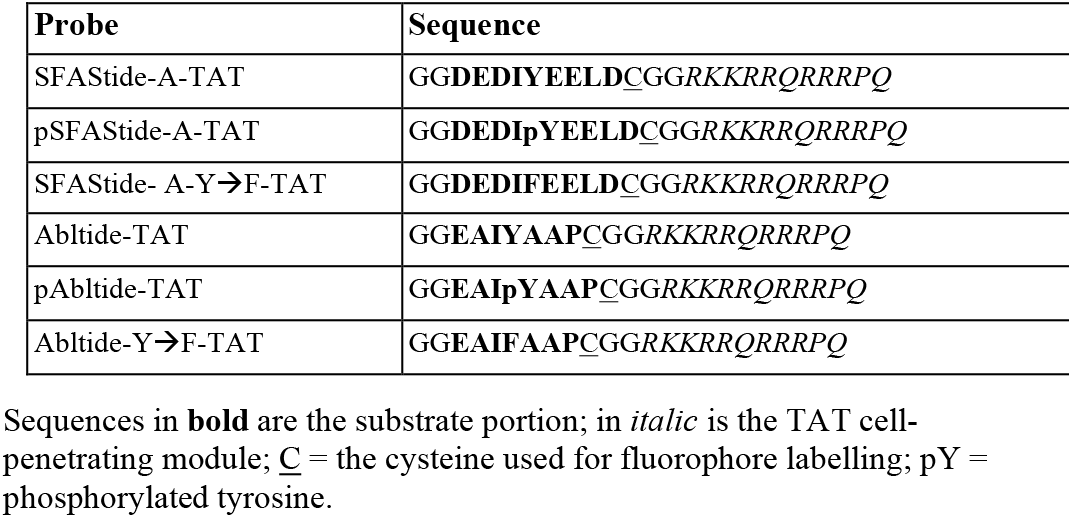
Peptide probe sequences

All three variants of SFAStide probes including the wild type (SrcF), synthetically phosphorylated (pSrcF), and Y→F (FmSrcF) mutants were labelled with DyLight 488 (DL488). These probes were tested in MDA-MB-231 cells for phosphorylation- and kinase activity-dependence of a fluorescence lifetime shift (Fig. 1, top and Fig. S1). The cells were incubated in media containing the probes (10 μM) for 2 hours. Imaging was performed as described in the Methods section (as in our previous publication).^1^

**Figure 1.**
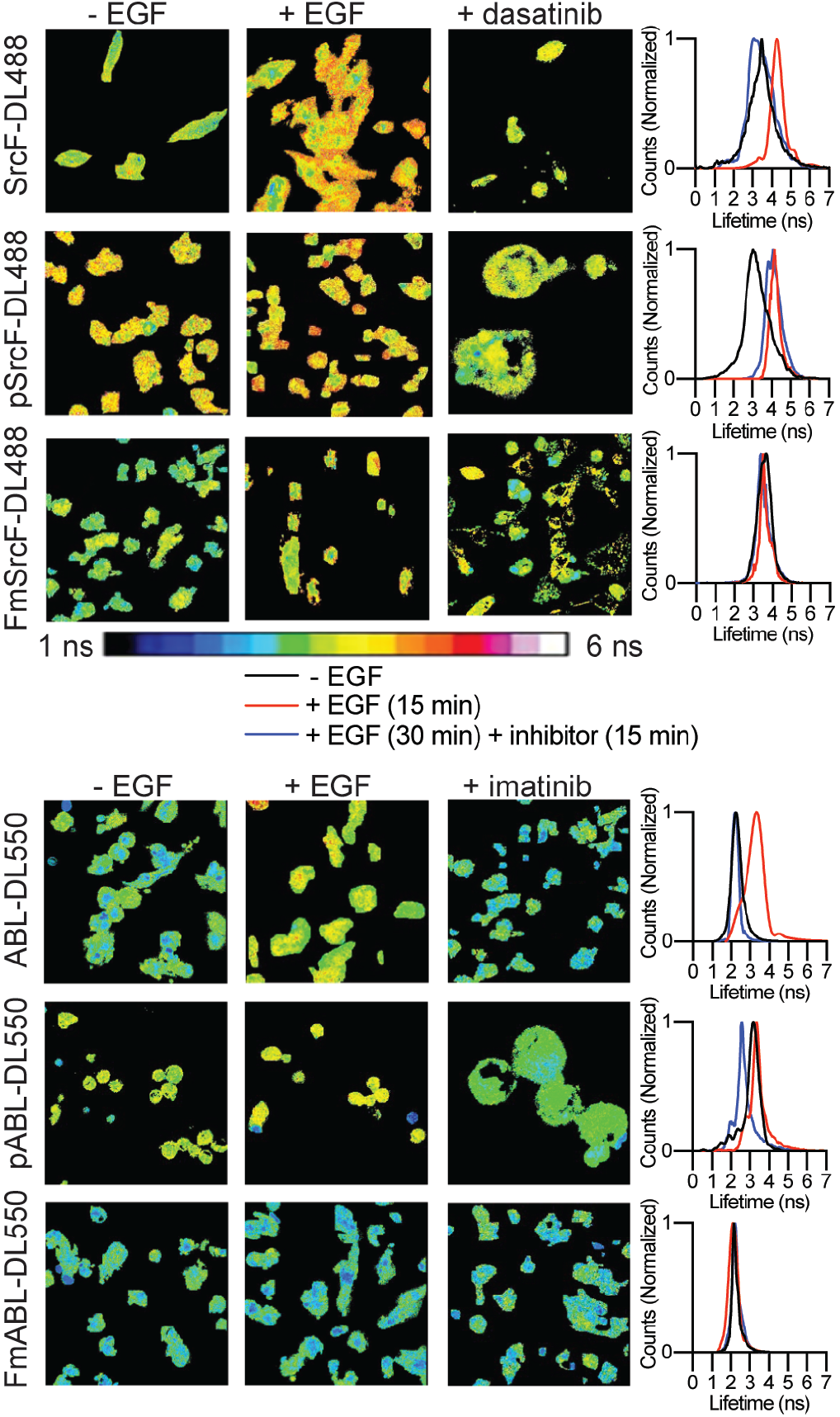
Probe characterization in MDA-MB-231 breast cancer cells for SrcF and Abl probes, labelled with DyLight488 or 550, respectively: SrcF-DL488, positive mutant (pSrcF-DL488) and negative Y->F mutant (FmSrcF-DL488) (top); ABL-DL550, positive mutant (pABL-DL550) and negative Y->F mutant (FmABL-DL550) (bottom). Cells were serum starved overnight and incubated with probe (10 μM) for 2h. DL488 or DL550 lifetime was measured before (−EGF) and after (+ EGF) treating the cells with epidermal growth factor (EGF, 10 ng, 15 min) to activate Src and Abl kinases. Longer lifetimes were observed in the wild type probes following stimulation as well as both before and after stimulation in case of the positive mutant probes, with no increase in the negative mutant, indicating that the longer lifetime species were consistent with peptide phosphorylation. After 15 min of EGF treatment, the inhibitor dasatinib (for Src probes) or imatinib (for Abl probes) was added (1 μM), decreasing the longer lifetime species. Images normalized to the respective lifetime scales and average lifetime histograms per image corresponding to the wild type, positive mutant and negative mutant probes for this experiment are shown.

We used a well-characterized model for Abl and Src family kinase activation downstream of epidermal growth factor receptor (EGFR) stimulation by EGF, with the dual Abl/Src-family inhibitor dasatinib to test inhibition.^8, 9^ As seen in Figure 1, longer lifetimes were observed for the wild type probes upon EGF stimulation, which can be attributed to phosphorylation by the respective kinase. Upon treatment with inhibitor, the lifetime distributions reverted to lower values and resembled distributions prior to stimulation indicating dephosphorylation of the probes by endogenous phosphatases. The Y→F mutants did not shift to longer lifetimes even with EGF stimulation, whereas the pY positive controls showed longer lifetimes preceding stimulation as well as following EGF treatment. Inhibitor treatment did result in lower lifetimes observed, consistent with phosphatase activity on the synthetically phosphorylated material not being balanced by corresponding kinase activity as we previously described.^1^ Abltide probes1 labelled with DyLight 488 (Fig. S3) (ABL-DL488) and DyLight 550 (ABL-DL550) (Fig. 1, bottom and Fig. S2) were similarly characterized. EGF stimulation was also used to activate the Abl kinase followed by treatment with the Abl inhibitor Imatinib. Their lifetime distributions showed the expected trend with activity stimulation and inhibition, indicating that regardless of the type of fluorophore (DyLight 488 or DyLight 550, or Cy5 as shown in our previous work^1^), all probes showed a phosphorylation dependent lifetime shift, despite intrinsically different lifetimes of the fluorophores.

As a preliminary application of the Abl and Src-family probes in multiplexed analysis of kinase activation and inhibition, we performed experiments to monitor the difference between the effects of dasatinib vs. imatinib on Abl and Src-family kinase activation in EGF-stimulated MDA-MB-231 cells (Fig. 2). For this purpose, we used SrcF-DL488 and ABL-DL550. Serum-starved cells were incubated in a 50:50 mixture of the two probes (10 μM each) followed by activation of both Src and Abl via EGF stimulation, and subsequent treatment with dasatinib or imatinib. Lifetime information from the SrcF-DL488 and ABL-DL550 probes was extracted using two detection channels per image. A threshold filter was applied to analyse only pixels with photon counts of at least 100 for the biexponential fit analysis. As a result, signal for each channel was not always equivalent (Fig. 2), giving rise to a punctate like appearance for the probe in some representative examples (Fig. S5)—however, this heterogeneity of intensity from experiment to experiment did not affect the consistency of lifetime distributions observed between replicates, highlighting the advantages of using lifetime as a probe read-out rather than intensity. Both probes exhibited longer lifetime upon EGF stimulation that was subsequently decreased by dasatinib (which inhibits Src family kinases and Abl). In contrast, if the cells were treated with imatinib (which only inhibits Abl, but not Src family kinases), only the ABL-DL550 probe show decreased lifetime while lifetime distribution of the SrcF-DL488 probe remains virtually unchanged (Fig. 2). These results show that the FLIM probes can be efficiently multiplexed by tagging with fluorophores of different colours characterized by distinct excitation wavelengths and emission bandwidths. They also demonstrate the utility of this approach for measuring kinase inhibitor selectivity in live cells.

**Figure 2:**
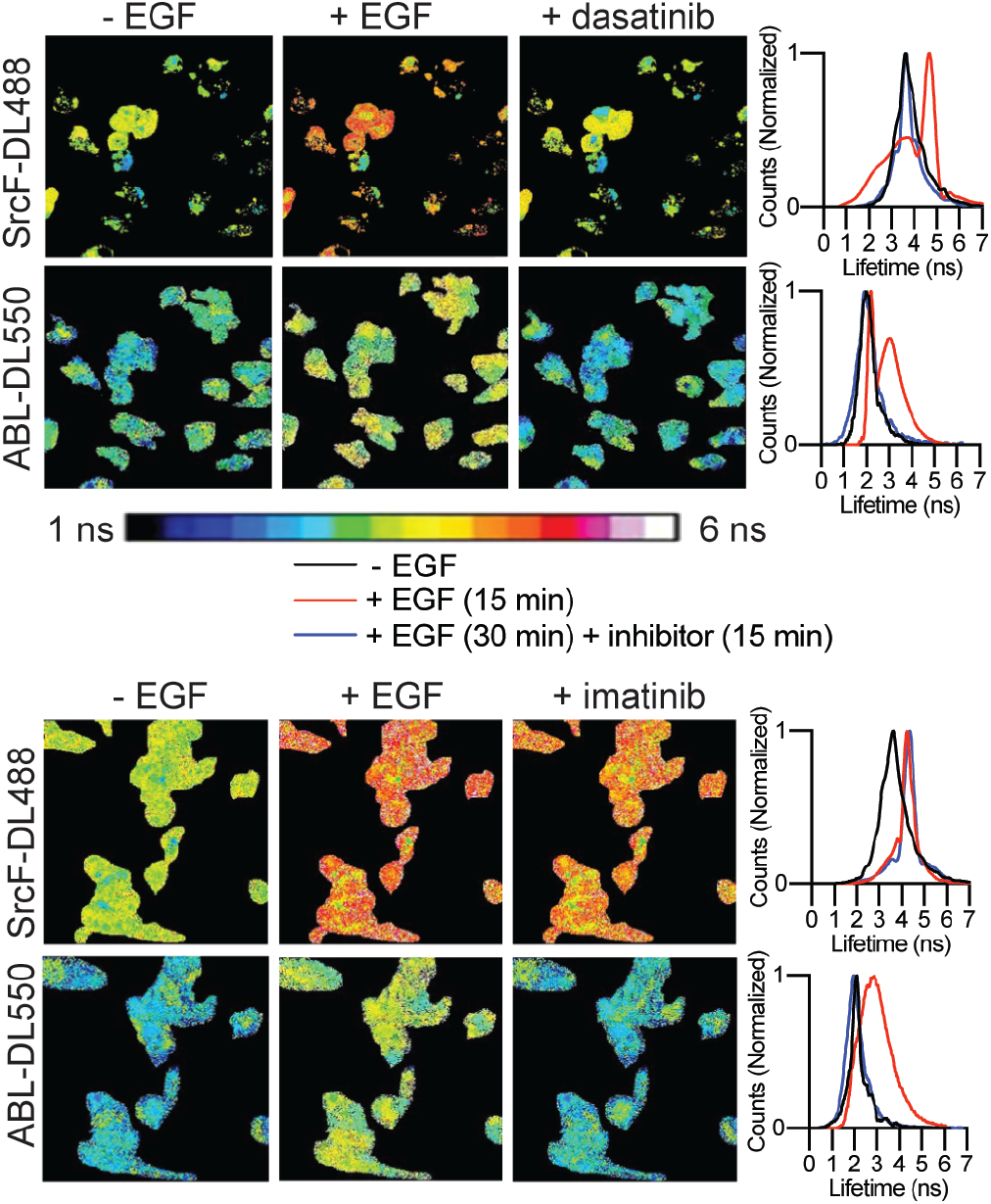
Dasatinib inhibits phosphorylation of both ABLtide and SFAStide probes (Top). Imatinib inhibits phosphorylation of ABLtide probe but not the SFAStide probe (Bottom). ABL-DL550 and SrcF-DL488 probes (10 μM each) were incubated with serum-starved MDA-MB-231 cells for 2 h. Kinase activity was stimulated with EGF (10 ng) for 15 min, followed by addition of inhibitor imatinib or dasatinib (1 μM) for an additional 15 min. Images normalized to the lifetime scale and average lifetime histograms corresponding to the green (top row) and red (bottom row) channels are shown.

Next, to understand the dynamics of phosphorylation of the probes by the respective kinases, the ABL-DL488 probe was imaged over a time course following stimulation with EGF and subsequent treatment with imatinib (Fig. 3). To quantify the relative amount of the longer lifetime species, we used a multi-exponential fitting model to extract the proportion of the longer vs. shorter lifetime species on a per-image basis from the time course experiments. This was done by fitting the averaged decay curve for each image to a bi-exponential model function by employing the Levenberg–Marquardt routine for non-linear least squares fitting,^10^ as further described in the Supporting Information. This non-biased fitting resolved two different lifetimes, as well as their fraction, to identify the average lifetime for each species. The relative intensity represented by the longer vs. the shorter lifetime species was then calculated and expressed as I_long_/I_total_. To extract the fraction of the longer lifetime component for individual pixels, a similar fitting analysis was performed on all pixels, iteratively, and the respective I_long_/I_total_ corresponding to each pixel was spatially mapped. The fraction of the longer lifetime species was mapped on a pixel by pixel basis in the representative FLIM images. The overall fraction of the longer lifetime component (averaged across all pixels) is plotted as at 5 min intervals (Fig. 3). The longer lifetime component (phosphorylated population) progressively increased following EGF treatment and reached steady state after 20 minutes, then steadily declined following imatinib treatment, stabilizing at a proportion similar to that at the beginning of the stimulation after ~20 minutes. Following 3-4 rounds of imaging, a gradual decrease in overall emission intensity was observed during time course experiments, due to photobleaching of the fluorophores (Fig. S6), however this did not affect lifetime measurements—again highlighting the advantages of lifetime as a read-out, especially for a time-course experiment.

**Figure 3:**
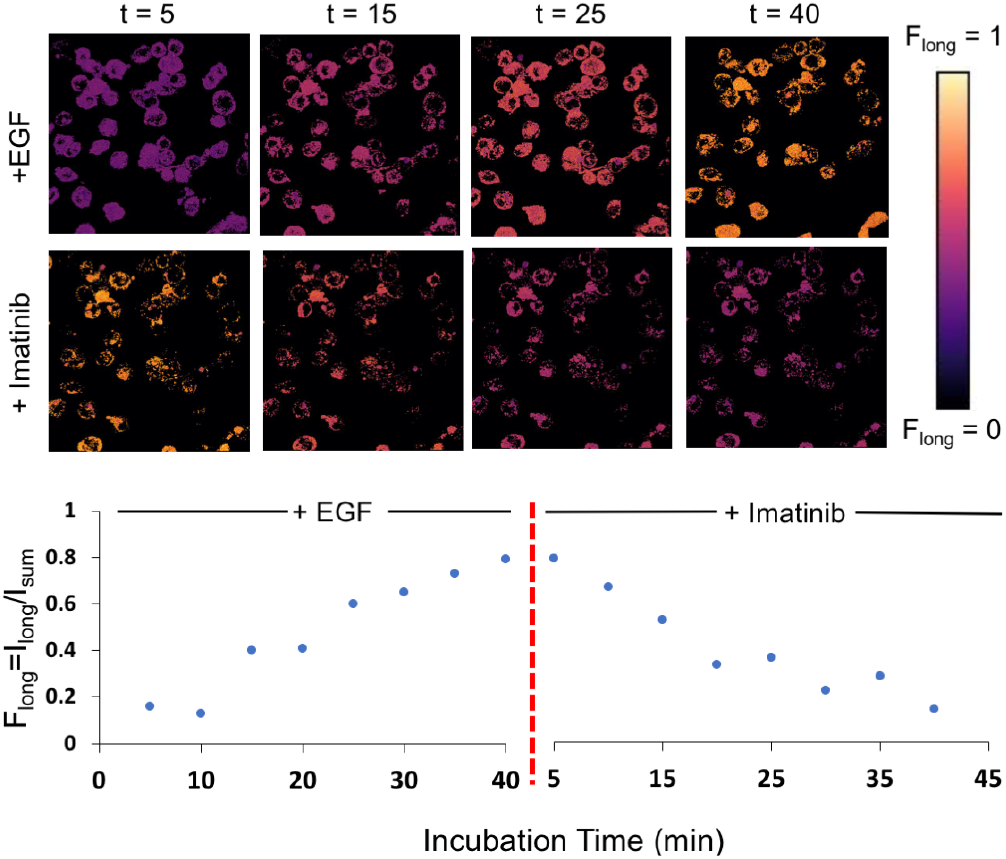
Quantitation of relative probe phosphorylation. FLIM data from timecourse experiments with ABL-DL488 probe were analysed via bi-exponential fitting as described in the supporting information, quantifying relative amount of phosphorylated probe per pixel following EGF treatment (top row) and subsequent imatinib treatment (bottom row). The relative fraction of phosphorylated probes (F_long_) is plotted as quantitative subcellular maps. The F_long_ value averaged over all the pixels is also plotted as a function of time.

Multiplexed time-course imaging was also performed, using Pulsed Interleaved Excitation (PIE), in which the system alternated between the two lasers (485 nm and 560 nm) and their respective detection channels in order to avoid excitation bleed through between the two fluorophores. The serum-starved MDA-MB-231 cells were incubated in a 50:50 mixture of SrcF-DL488 and Abl-DL550 (10 μM each), excess probe was removed and replaced with fresh serum-free colourless media, and the cells were imaged at 5 min intervals after treating with EGF (40 min) and dasatinib (an additional 40 min). The green and red channels were separately analysed to obtain lifetime information corresponding to SrcF-DL488 (Fig. 4A) and ABL-DL550 (Fig. 4B), respectively. As expected, lifetimes for both probes increased over time after EGF stimulation and decreased after adding dasatinib.

**Figure 4:**
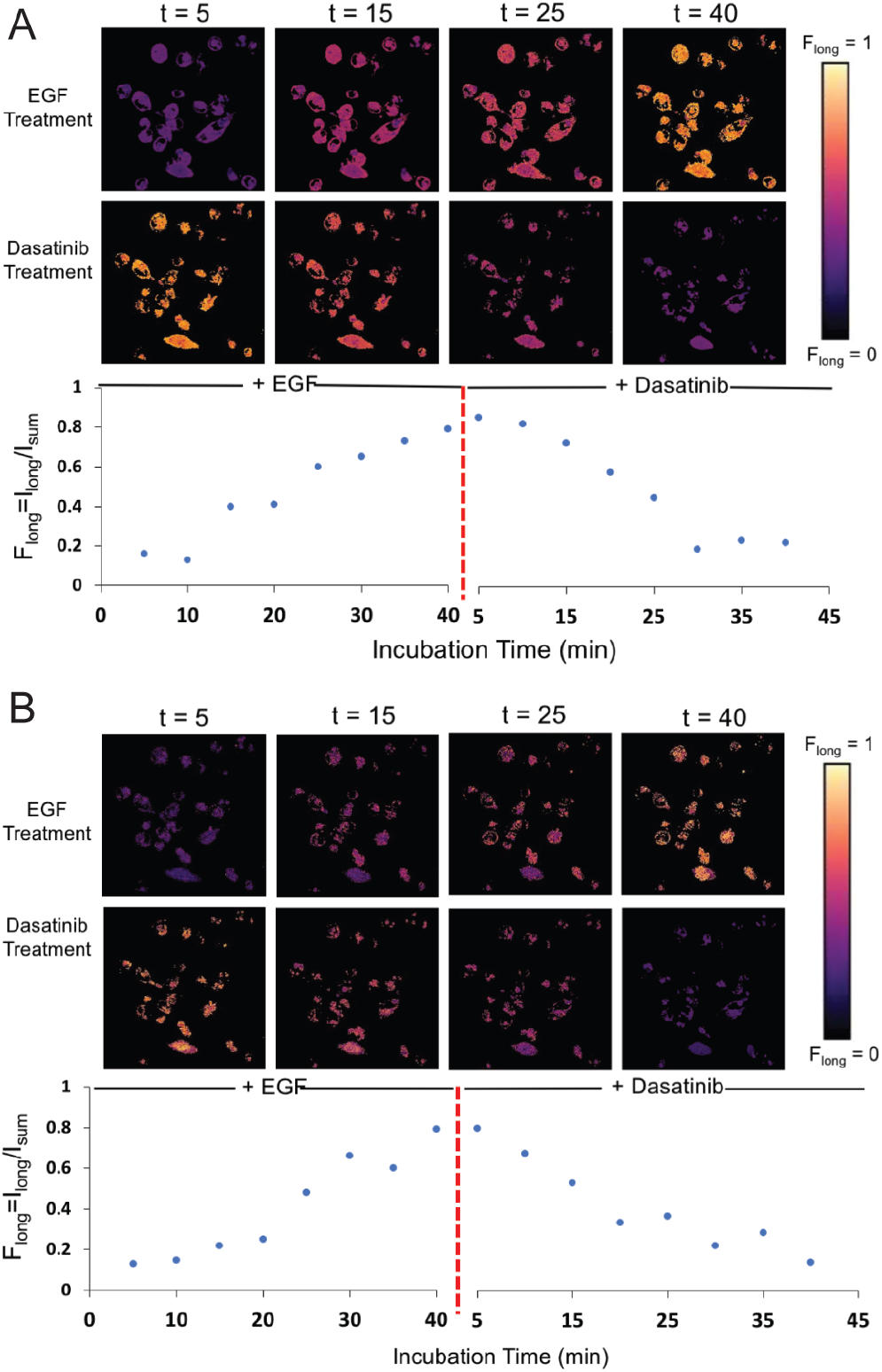
Quantitation of relative probe phosphorylation in multiplexed probes. FLIM data from EGF stimulation and imatinib inhibition experiments of (A) SrcF-DL488 and (B) ABL-DL550 probes were analysed via bi-exponential fitting to quantify the relative amount of phosphorylated probe per pixel, following EGF treatment (top row) and imatinib treatment (bottom row). As in Fig. 4, the relative fraction of phosphorylated probes (F_long_) is plotted as quantitative subcellular maps. The F_long_ value averaged over all the pixels is also plotted as a function of time.

These analyses illustrate the potential to obtain subcellular maps of FLIM probe phosphorylation that could ultimately be used for more refined quantification of kinase activities in specific areas of the cell, paving the way towards incorporation of FLIM probe data into complex models of signalling behaviour both between and within individual cells.

## Conclusions

In this work, we showed that combining different fluorophores and peptides into Abl and Src-family kinase FLIM probes exhibited similar phosphorylation-dependent lifetime behaviour, implying that the effects of phosphorylation via kinase activity on fluorescence lifetime of these probes are likely to be broadly applicable. We also demonstrated two-color imaging of Abl and Src-family kinase activities downstream of EGFR stimulation by EGF, and were able to distinguish between subsequent inhibition of both kinases with the Abl/Src inhibitor dasatinib vs. only Abl kinase with imatinib. Finally, we demonstrated for the first time for this kind of probe that the relative proportions of the different lifetime species can be quantified on a pixel-by-pixel basis to infer relative probe phosphorylation. These results show the generality and multiplexability of this approach, demonstrate the potential to quantify proportions of phosphorylated and unphosphorylated species via multicomponent fitting, and provide a specific strategy for more advanced quantitative study of subcellular kinase activity using these and similar probes.

## ‡ Acknowledgements

Cells used in this work were a kind gift from Prof. Robert Geahlen (Purdue University Department of Medicinal Chemistry and Molecular Pharmacology). This work was supported by the Purdue Center for Cancer Research (NPD and JI) and the NCI/NIH (R01CA182543 and R33CA217780 to LLP).

Electronic Supplementary Information (ESI) available: Includes detailed experimental and data analysis procedures, probe characterization data, and additional fluorescence images. See biorXiv entry.

