## Supplemental information for "Multiplexable fluorescence lifetime imaging (FLIM) probes for Abl and Src-family kinases"

#### *Experimental procedures*

**Peptide Biosensors.** Peptide sensors were previously designed and developed by the Parker lab[1, 2]. Complete sequences are available in Table 1. Briefly, Fmoc solid phase peptide chemistry was used to synthesize each sensor, which consists of recognition sequence modified with cysteine as reporter anchoring site, fluorescence reporter and Tat penetrating sequence to aid cellular delivery. The fluorophore maleimides (DyLight 550 or 488 from ThermoFisher Scientific) were conjugated to the cysteines for each respective sensor in buffer containing 6M guanidinium-HCl, 100 mM phosphate, and 10 mM TCEP, at pH 6.5, checking desalted aliquots with MALDI-TOF MS until reaction had gone to completion (typically ~1-2 h). Sensor peptides were purified using HPLC (Agilent 1200 preparative system, Zorbax C18 column, 20 mm inner diameter, 250 mm length). Final labeled peptides were characterized with LC/MS (see pp. 12-15).

**Cell Culture.** Human MDA-MB-231 were purchased from ATCC. Cells were cultured in DMEM medium supplemented with 10% FBS and 1% Penicillin/Streptomycin at 37°C, 5% $CO_2$.

**Sensor Delivery and Imaging.** Cells were serum starved for 24 h prior to EGF stimulation experiments. Live cells were then incubated with the respective biosensors (10 μM) in serum free medium for 45 minutes prior to FLIM imaging experiments. Cells were washed with PBS and maintained in the low background fluorescence medium, FluoroBrite DMEM (Gibco, USA) in a top incubator live cell imaging chamber (TOKAI HIT INUBG2ASFP). The live cell chamber was equipped with temperature feedback control to maintain stable temperature at 37$^{o}$C.

**TCSPC-FLIM.** FLIM experiments utilized a Nikon A1R laser scanning confocal microscope equipped with a FLIM, FRET, and FCS enabled system (LSM Upgrade Kit, PicoQuant, Germany)[3-9] that allows precise recording of the arrival time of photons at the detector. The Instrument Response Function (IRF)[3] was taken into consideration during measurement and data analysis. Two picosecond pulsed laser lines 488 and 561 nm were used as excitation sources and the emitted light was guided through a 50 µm pinhole. Two single photon avalanche diode detectors ( PDM SPAD, PMA Hybrid PMT.) were used to collect emitted photons. A 520±30 nm emission filter was placed in front of the SPAD to ensure collection from DL488 while a 580±40 nm filter was placed in front of the Hybrid PMT to collect emission from the Dylight 550 fluorophore. Each photon was tagged with a time stamp that signified its arrival time in the detector after a laser pulse, using the time correlated single photon counting (TCSPC) in the Time Tagged Time Resolved Single Photon Mode (TTTR) (Picoharp 300, Picoquant). Average fluorescence lifetimes were measured and displayed as color contrast unless otherwise specified, using the color scales indicated for each image. For quantitative analysis, lifetime decay per image was fitted per pixel with bi-exponential fitting employing an iterative Levenberg-Marquadt algorithm (Equation 1).

$I(t) = \sum_{i=1}^{n} \alpha i exp(-t/\tau i) + C$ ........................(Equation 1)

where I(t) is the fluorescence intensity at time t after the excitation pulse, n is the total number of decay components in the exponential sum, and C is a constant pertaining to the level of background light noise. The variables $\tau$i and $\alpha$i are the fluorescence lifetime and fractional contribution of the emitting species, respectively.

**Bi-exponential fitting**

I(t) = $\alpha 1 \exp\left(-\frac{t}{\tau 1}\right) + \alpha 2 \exp\left(-\frac{t}{\tau 2}\right) + C$ … … . ( Equation 2)

To ensure unbiased fitting, fluorescence decay curves collected in each pixel were fitted with (Equation 2) until two criteria were met: 1. sufficient Chi value (<1.3) and 2. no residual pattern was observed. Fitting resulted in two fluorescence lifetime values ($\tau 1$ and $\tau 2$) as well as their population ratio in each pixel ($\alpha 1$ *and* $\alpha 2$) . Fitting was performed in SymphoTime (Picoquant) software. The results were exported to MATLAB to separate matrices of lifetime and matrices of intensity of each component. The total ratio of matrices describing the fractional intensity of longer lifetime over total intensity was calculated in MATLAB. The matrices were then exported to ImageJ for mapping. A schematic of the instrumentation and analysis steps are provided below.

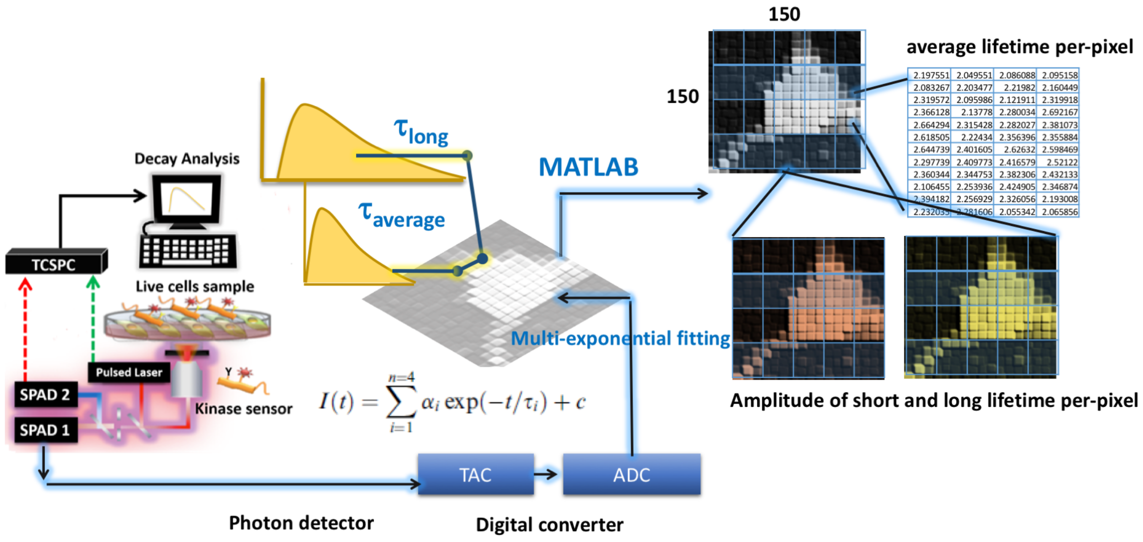


**Analysis schematic.** Cells are imaged using a TCSPC FLIM microscope, and files are analyzed by deconvoluting multiple lifetimes via multi-exponential fitting. Proportional amplitudes for different lifetime components can be extracted and re-plotted as fractional intensity of longer lifetime species.

### *Additional microscopy images*

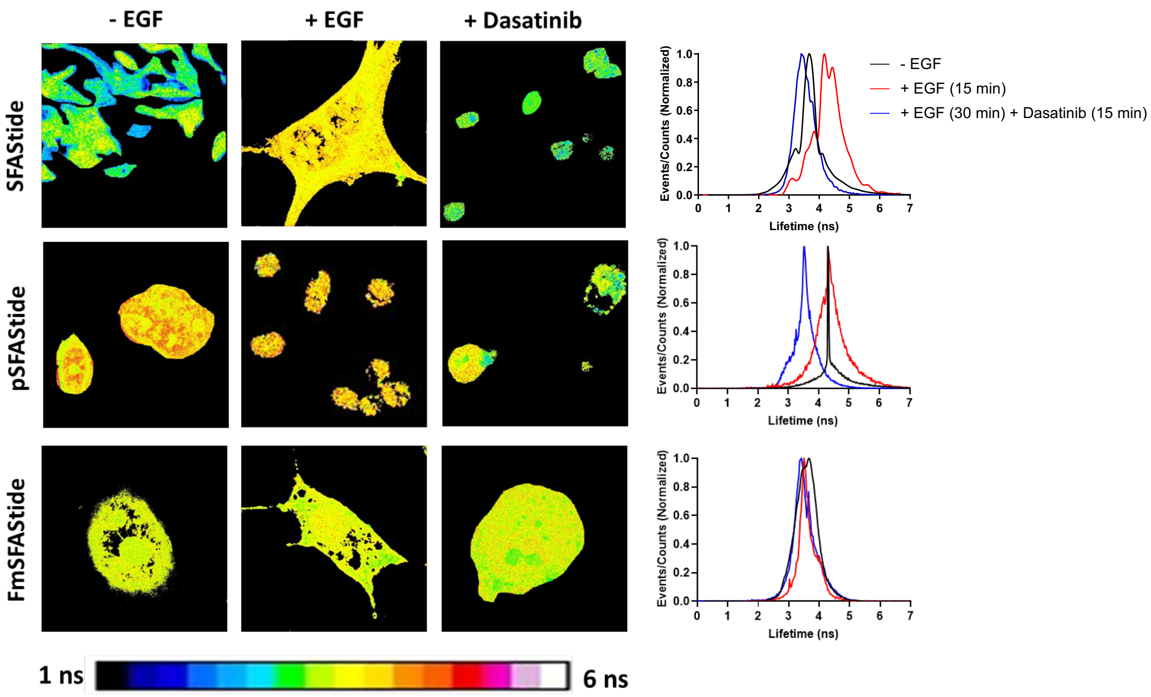


**Figure S1. SFAS-DL488 probes analyzed in MDA-MB-231 cells (duplicate experiment from Figure 1 of main manuscript).** The probes were synthesized and labeled with DyLight488 maleimide. MDA-MB-231 cells were incubated with probes and imaged before treatment, following EGF treatment and following inhibitor treatment. Representative lifetime mapped images and lifetime histogram plots are shown.

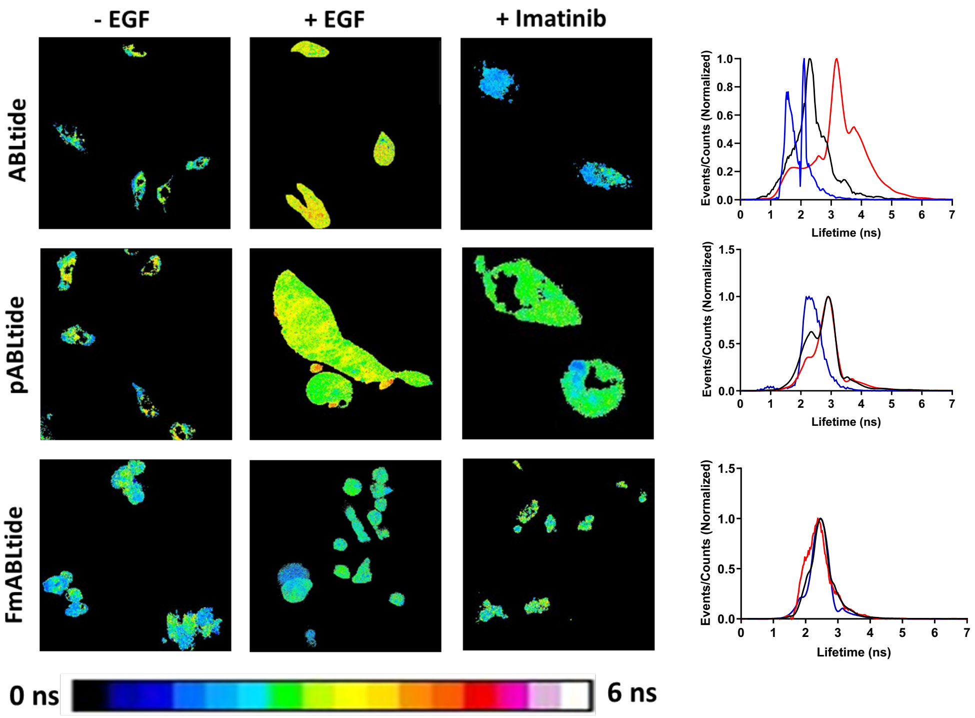


**Figure S2. Abl-DL550 probes analyzed in MDA-MB-231 cells (duplicate experiment from Figure 1 of main manuscript).** The probes were synthesized and labeled with DyLight550 maleimide. MDA-MB-231 cells were incubated with probes and imaged before treatment, following EGF treatment and following inhibitor treatment. Representative lifetime mapped images and lifetime histogram plots are shown.

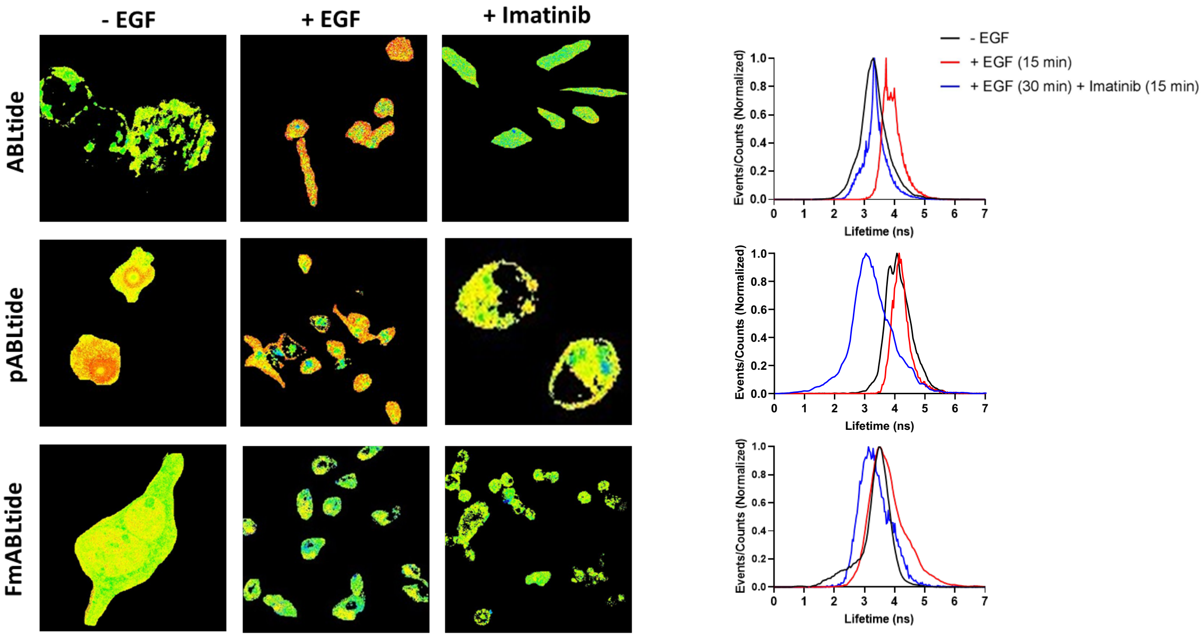


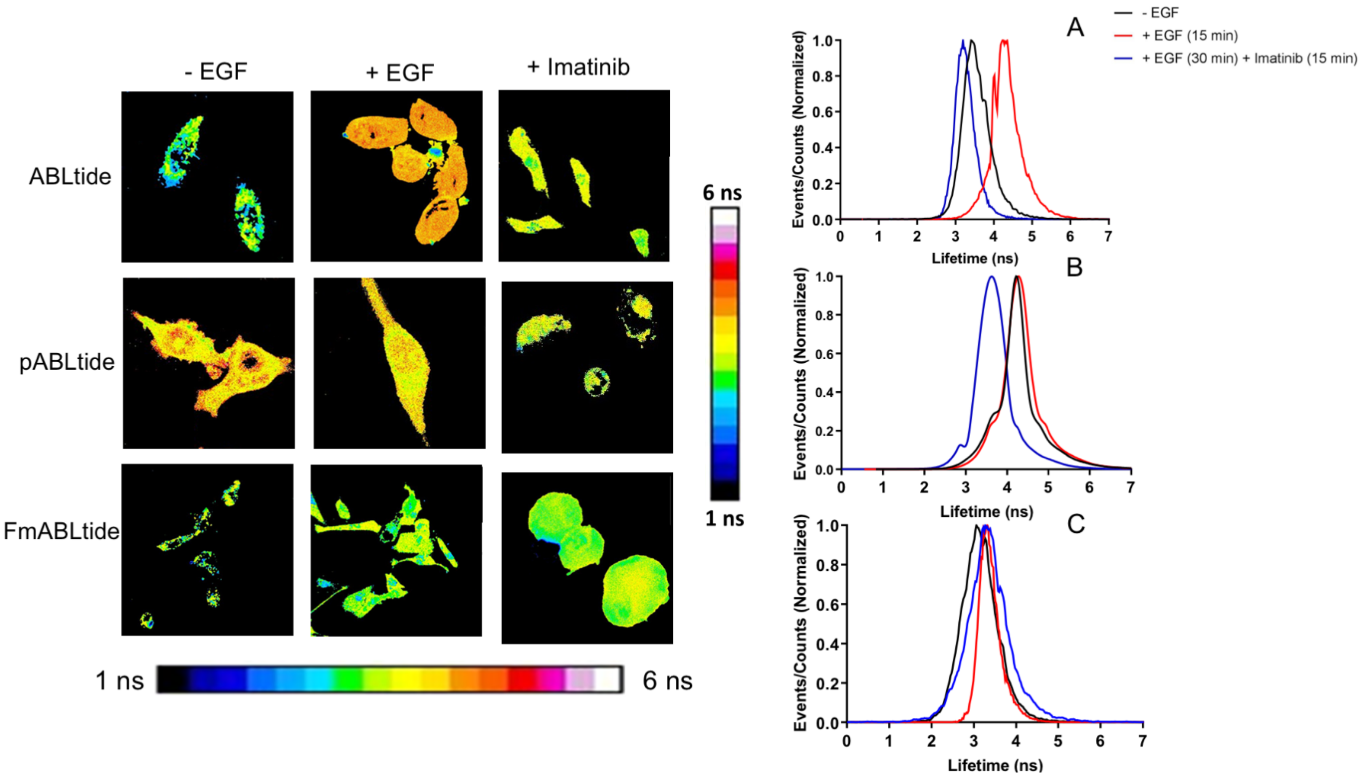


**Figure S3. Abl-DL488 probes analyzed in MDA-MB-231 cells.** The probes were synthesized and labeled with DyLight488 maleimide. MDA-MB-231 cells were incubated with probes and imaged before treatment, following EGF treatment and following inhibitor treatment. Representative lifetime mapped images and lifetime histogram plots are shown.

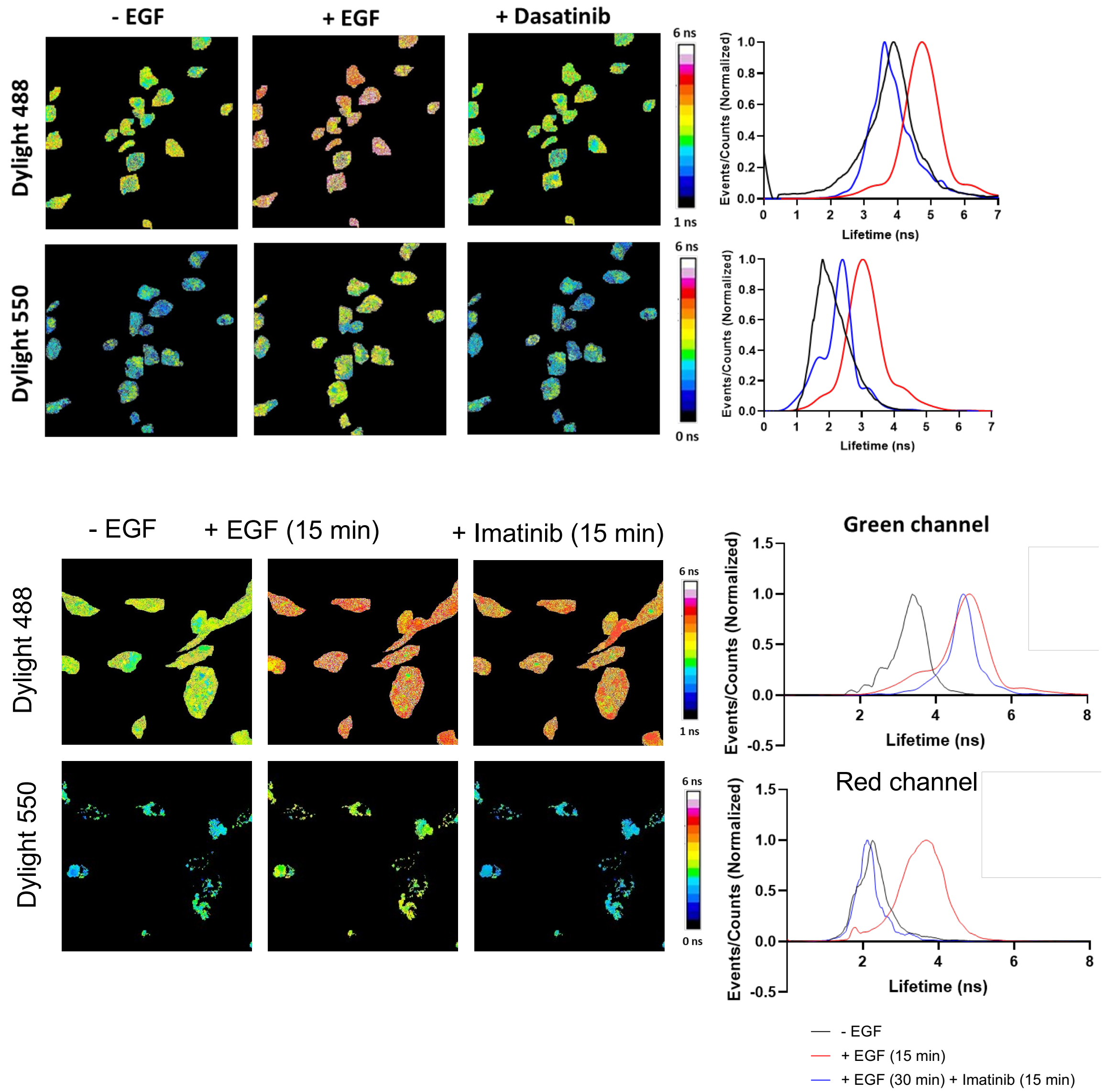


**Figure S4. Multiplexing of SFAS-DL488 and ABL-DL550 probes in MDA-MB-231 cells.** Representative FLIM images and lifetime histograms from duplicate experiments performed for multiplexed analysis in Figure 2 of the main manuscript.

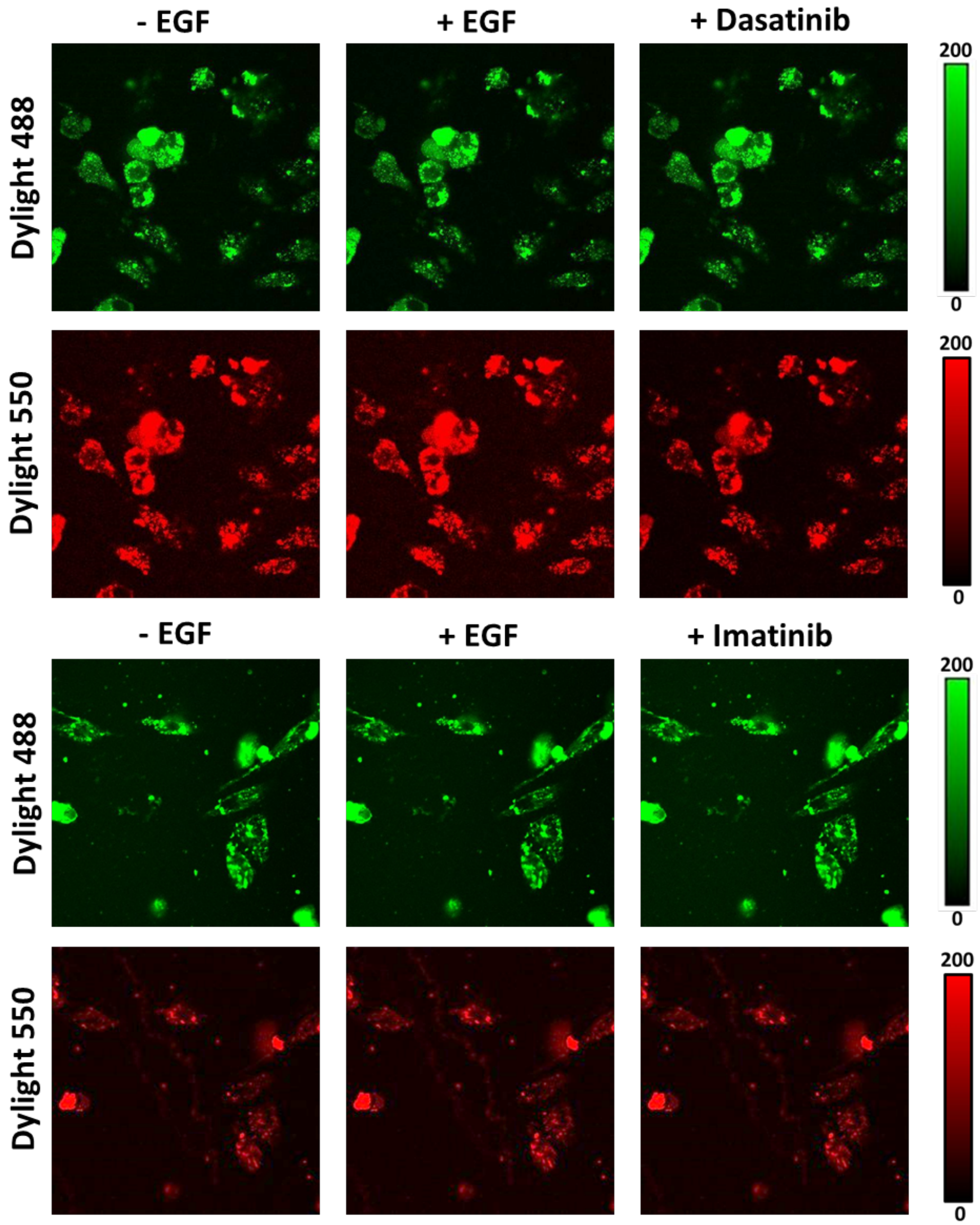


**Figure S5. Fluorescence Intensity images of SFAS-DL488 (green channel) and Abl-DL550 probes (red channel) from multiplexed analysis.** Top: Dasatinib treatment from Fig. 2 of the main manuscript. Bottom: Imatinib treatment from duplicate experiment shown in Fig. S4.

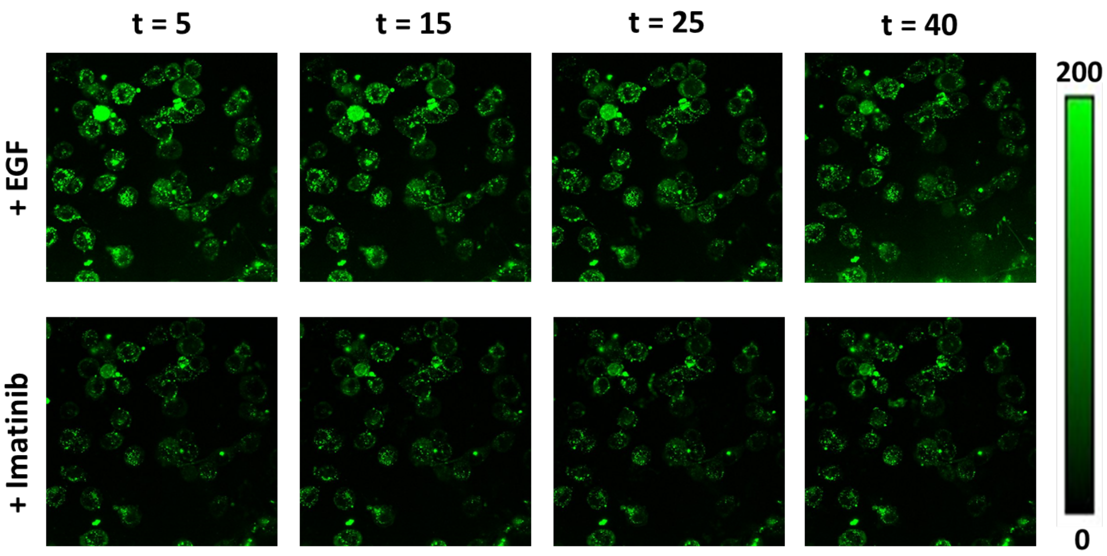


**Figure S6. Fluorescence Intensity images of ABL-DL488 probes from time course analysis in Figure 3 of the main manuscript.**

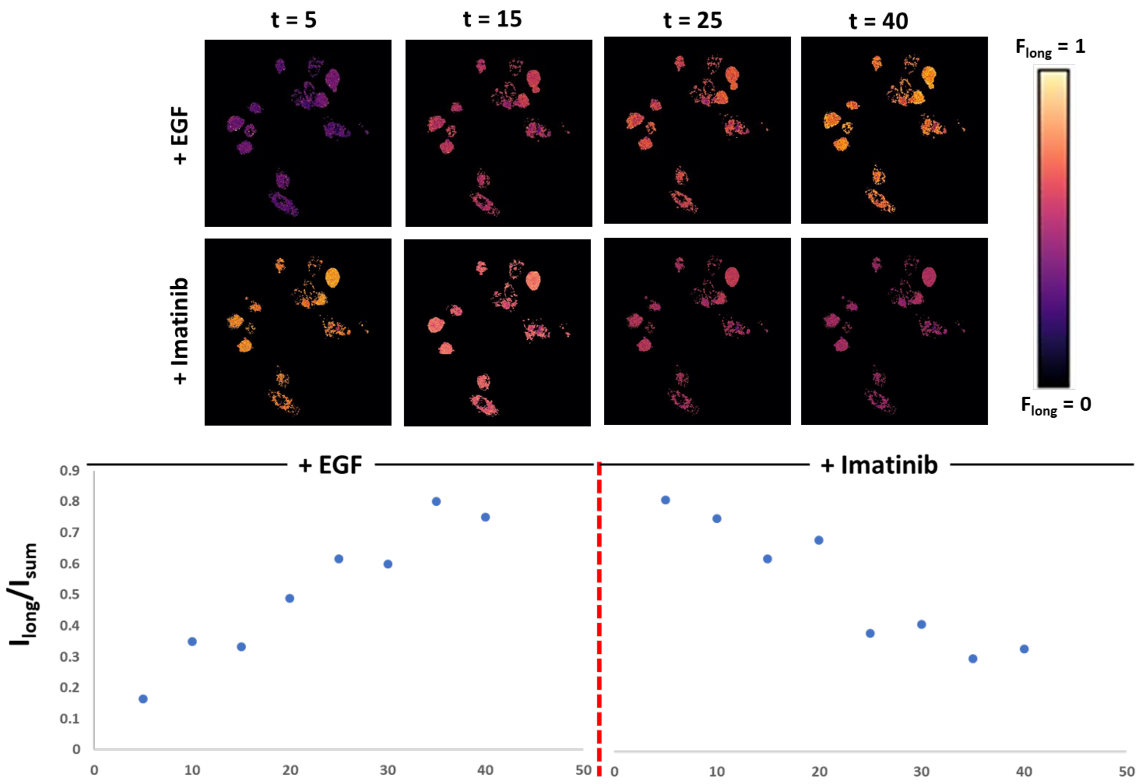


**Figure S7. Relative fraction of phosphorylated ABL-DL488 probes spatially mapped in a pixel-pixel basis and average value plotted as a function of time from a duplicate time course experiment in Figure 3 of the main manuscript.**

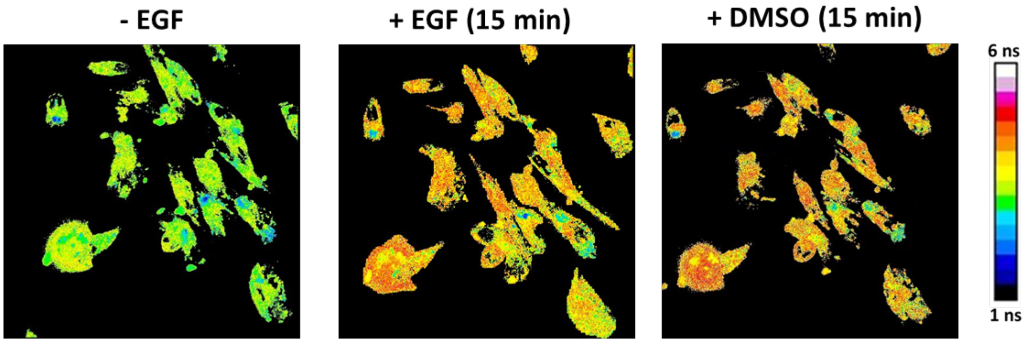


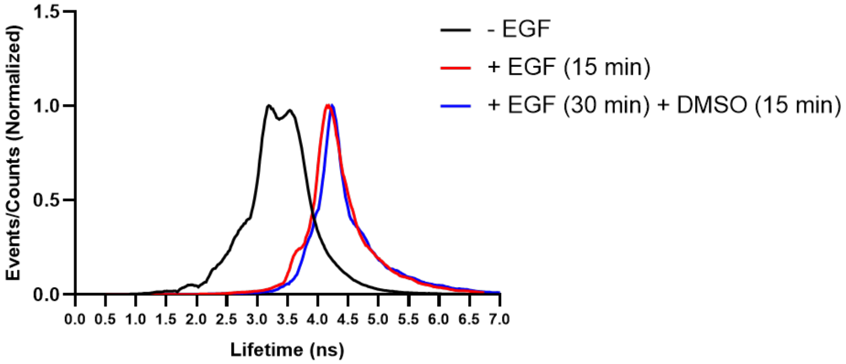


**Figure S8. DMSO Control.** MDA-MB-231 cells were incubated with Abl-DL488 probes and imaged before treatment, following EGF treatment and DMSO treatment. Representative lifetime mapped images and lifetime histogram plots are shown.

### *Peptide characterization data*

Table S1. Peptides used in this work

| **Peptide** | **Sequence** | **Characterization data available?** | **Fluorophore label** |
|---|---|---|---|
| SFAS-DL488 | GG**DEDIYEELDC**$_{\text{DyLight488}}$GG*RKKRRQRRRPQ* | Yes (HPLC) | DyLight 488 maleimide |
| pSFAS-DL488 | GG**DEDIYEELDC**$_{\text{DyLight488}}$GG*RKKRRQRRRPQ* | Yes (HPLC) | DyLight 488 maleimide |
| FmSFAS-DL488 | GG**DEDIFEELDC**$_{\text{DyLight488}}$GG*RKKRRQRRRPQ* | Yes (HPLC) | DyLight 488 maleimide |
| Abl-DL488 | GG**EAIYAAPC**$_{\text{DyLight488}}$GG*RKKRRQRRRPQ* | Yes (HPLC) | DyLight 488 maleimide e |
| pAbl-DL488 | GG**EAIpYAAPC**$_{\text{DyLight488}}$GG*RKKRRQRRRPQ* | Yes (HPLC) | DyLight 488 maleimide |
| FmAbl-DL488 | GG**EAIFAAPC**$_{\text{DyLight488}}$GG*RKKRRQRRRPQ* | Yes (HPLC) | DyLight 488 maleimide |
| Abl-DL550 | GG**EAIYAAPC**$_{\text{DyLight550}}$GG*RKKRRQRRRPQ* | Yes (HPLC) | DyLight 550 maleimide |
| Abl-DL550 | GG**EAIYAAPC**$_{\text{DyLight550}}$GG*RKKRRQRRRPQ* | Yes (HPLC) | DyLight 550 maleimide |
| Abl-DL550 | GG**EAIYAAPC**$_{\text{DyLight550}}$GG*RKKRRQRRRPQ* | Yes (HPLC) | DyLight 550 maleimide |

SFAS-DL488 GGDEDIYEELDC$_{\text{DyLight488}}$GGRKKRRQRRRPQ MW 3796.3

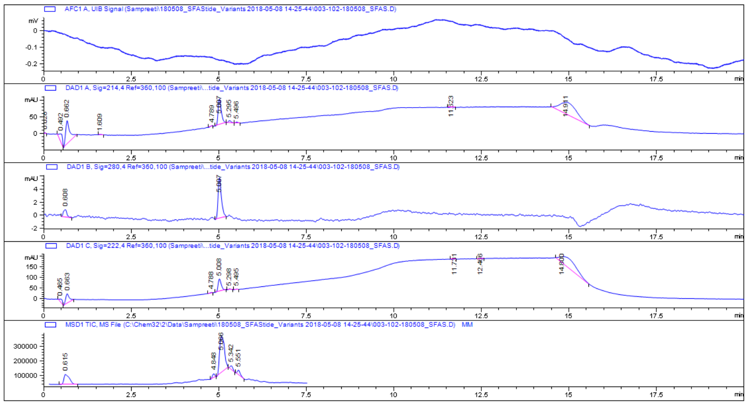

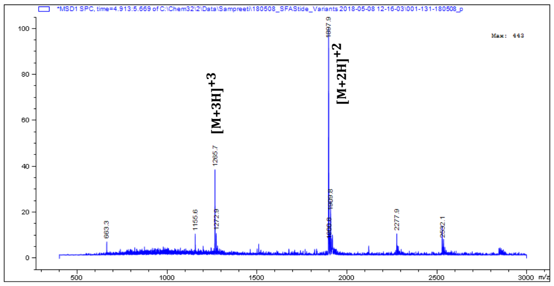

pSFAS-DL488 GGDEDIYEELDC$_{DyLight488}$GGRKKRRQRRRPQ MW 3874.3

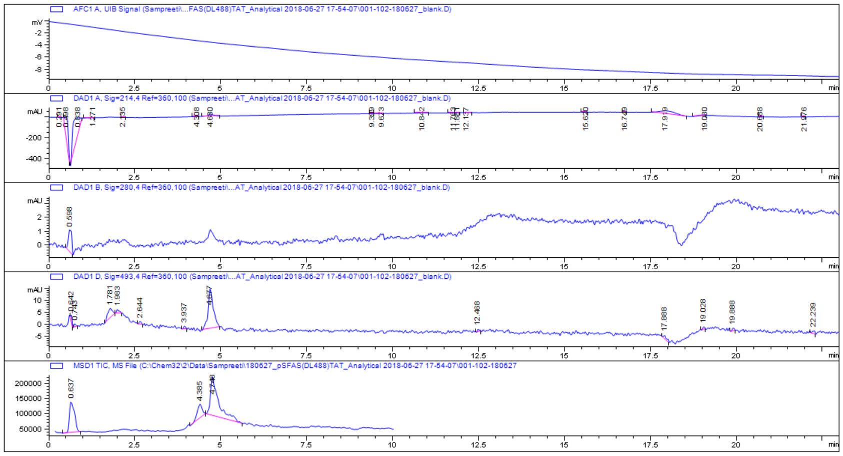

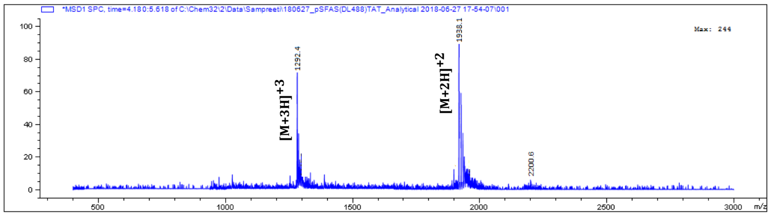


FmSFAS-DL488 GGDEDIFEELDC$_{DyLight488}$GGRKKRRQRRRPQ MW 3780.3

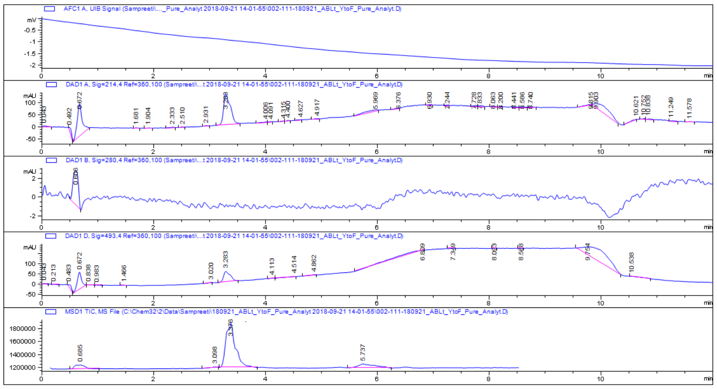

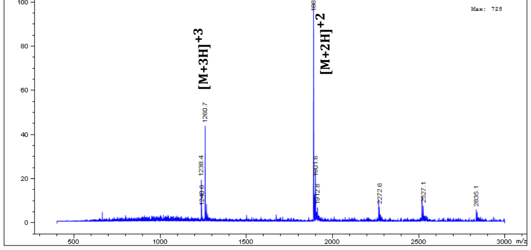

Abl-DL488 GG**EAIYAAP**<u>C</u>$_{\text{DyLight-488}}$GG*RKKRRQRRRPQ* MW = 3390.0

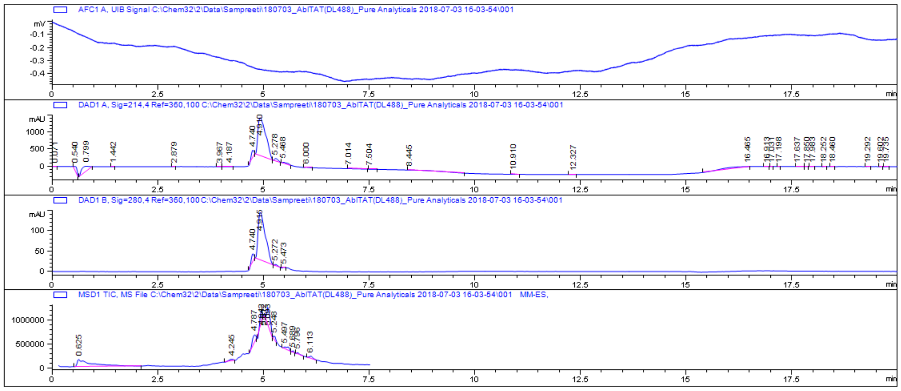


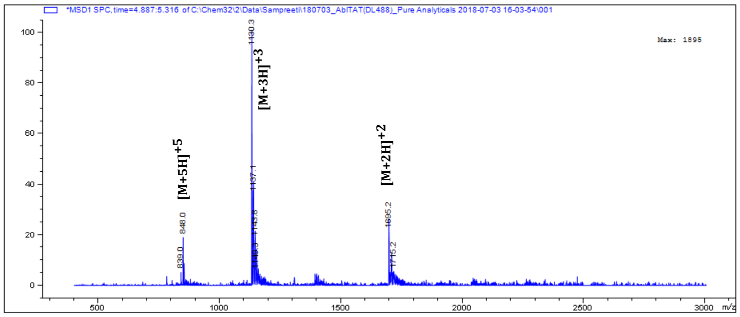


FmAbl-DL488 GG**EAIFAAP**<u>C</u>$_{\text{DyLight-488}}$GG*RKKRRQRRRPQ* MW = 3374.0

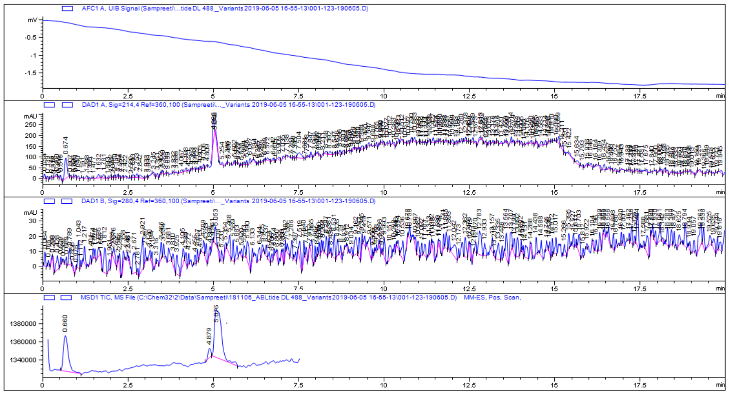


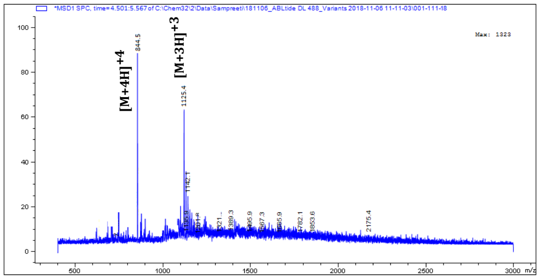

pAbl-DL488 GG**EAIYAAP**C$_{\text{DyLight-488}}$GG*RKKRRQRRRPQ* MW = 3468.0

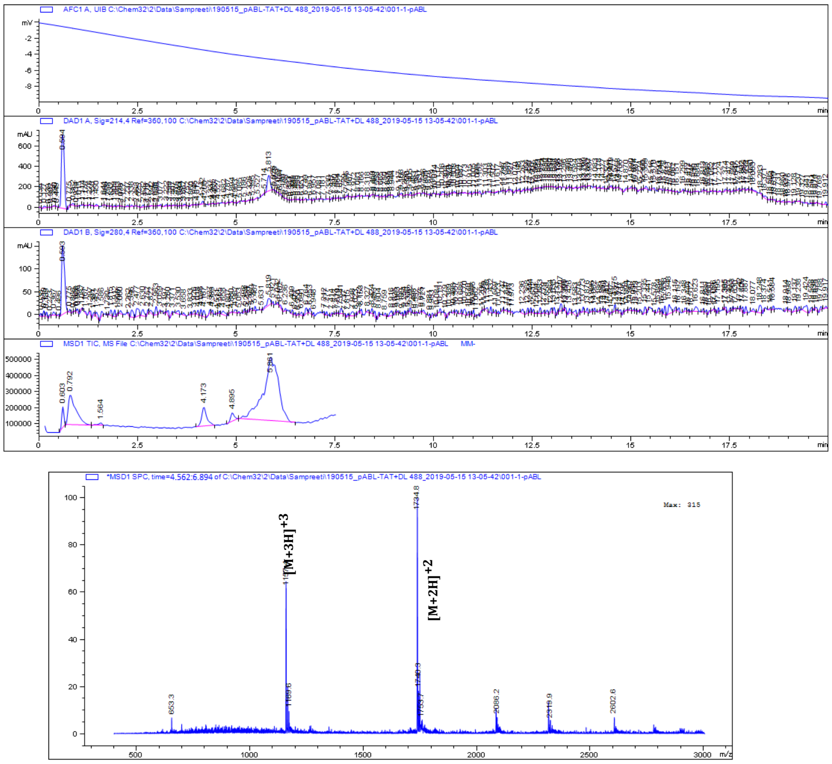


AbI-DL550 GG**EAIYAAP**C$_{\text{DyLight-550}}$GG*RKKRRQRRRPQ* MW = 3612.0

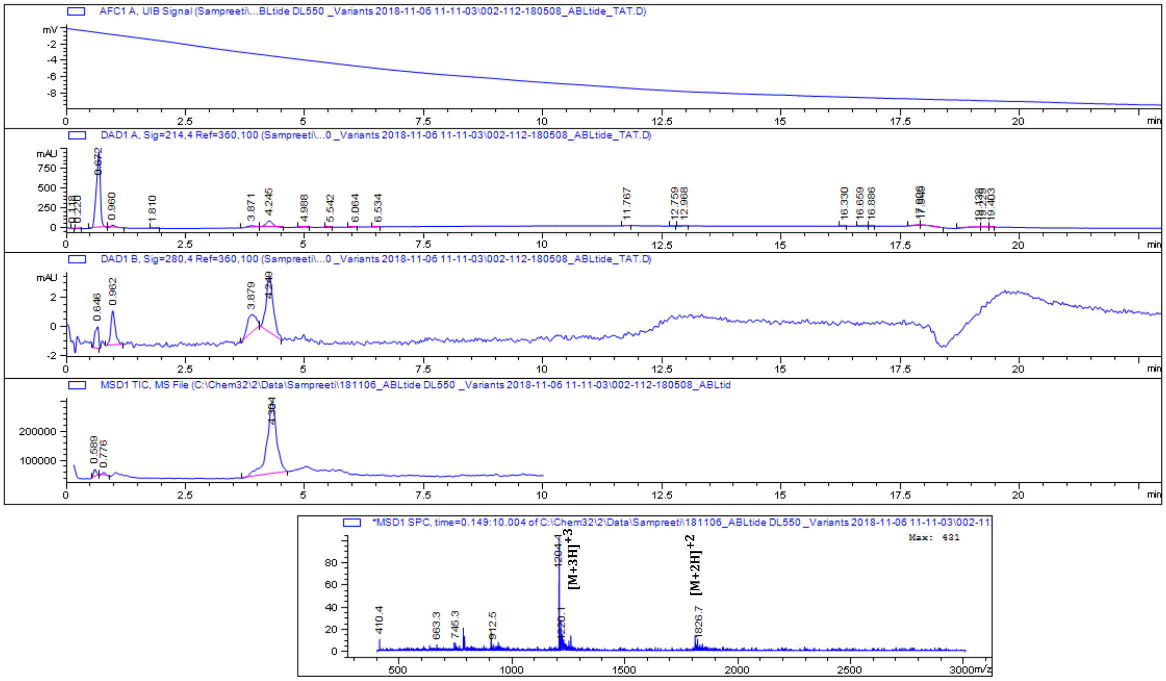

pAbl-DL550 GG**EAIpYAAP**$\underline{C}_{\text{DyLight-550}}$GG*RKKRRQRRRPQ* MW = 3690.0

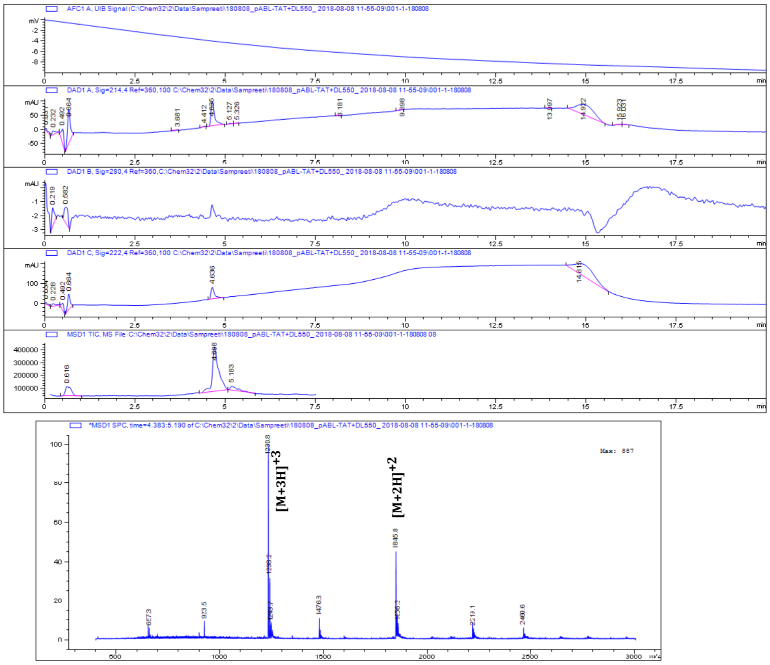


FmAbl-DL550 GG**EAIFAAP**$\underline{C}_{\text{DyLight-550}}$GG*RKKRRQRRRPQ* MW = 3596

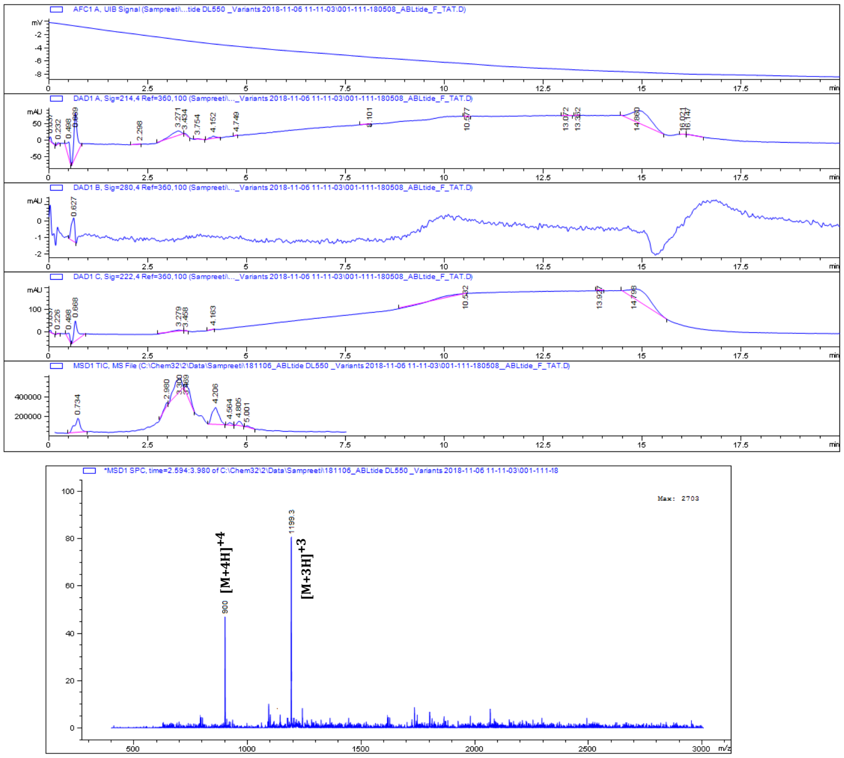
